# 24-Nor-ursodeoxycholic acid counteracts T_H_17/Treg imbalance and ameliorates intestinal inflammation by restricting glutaminolysis in differentiating T_H_17 cells

**DOI:** 10.1101/2022.02.10.479975

**Authors:** Ci Zhu, Nicole Boucheron, Ramona Rica, Valentina Stolz, Emina Halilbasic, Thierry Claudel, Osamah Al-Rubaye, Alexander Lercher, Maximilian Baumgartner, Lisa Sandner, Teresa Preglej, Marlis Alteneder, Veronika Mlitz, Claudia D. Fuchs, Daniela Hainberger, Jelena Remetic, Anna Ohradanova-Repic, Philipp Schatzlmaier, Tatjana Stojakovic, Hubert Scharnagl, Shinya Sakaguchi, Andreas Bergthaler, Hannes Stockinger, Wilfried Ellmeier, Michael Trauner

## Abstract

**Objective:** 24-Nor-ursodeoxycholic acid (NorUDCA) is a novel therapeutic bile acid for treating primary sclerosing cholangitis (PSC), an immune-mediated cholestatic liver disease. Since PSC strongly associates with inflammatory bowel diseases (IBD) driven by T_H_17/Treg imbalance, we aimed to explore NorUDCA’s immunomodulatory potential on intestinal T_H_17/Treg balance.

**Design:** NorUDCA’s impact on T_H_17/Treg tissue distribution was first assessed in *Mdr2*^*–/–*^ mouse model of PSC. We specifically investigated NorUDCA’s effect on modulating T_H_17/Treg balance in a CD4^+^ T cell driven colitis model induced by adoptive transfer of CD25^−^CD44^low^CD45RB^high^CD4^+^ T_Naïve_ cells into *Rag2*^*–/–*^ mice, mimicking human IBD. Mechanistic studies were performed using molecular approaches, flow cytometry and metabolic assays in murine T_H_17 cells *in vitro*. NorUDCA’s signaling effects observed in murine system were further validated in circulating CD4^+^ T cells from PSC patients with co-existing IBD.

**Results:** NorUDCA promoted Treg generation in both liver and intestine in the *Mdr2*^*–/–*^ model. In the experimental IBD model, NorUDCA attenuated intestinal immunopathology. Mechanistically, NorUDCA demonstrated strong immunomodulatory efficacy in counteracting T_H_17/Treg imbalance by restricting glutaminolysis in differentiating T_H_17 cells, thus suppressed α-Ketoglutarate-dependent mTORC1 activation, glycolysis and enhanced FOXP3 expression. NorUDCA’s impact on mTORC1 signaling was further confirmed in circulating CD4^+^ T-cells from PSC patients with IBD.

**Conclusion:** NorUDCA possesses direct immunometabolic modulatory potency to counteract T_H_17/Treg imbalance and ameliorate excessive T_H_17 cell driven intestinal immunopathology. These findings extend future clinical applications of NorUDCA for treatment of T_H_17 cell-mediated disorders along the gut-liver axis and beyond.

**Significance of this study:** *What is already known on this subject?:*

- PSC is an immune-mediated cholestatic liver disease highly associated with IBD where T_H_17/Treg imbalance drives immunopathogenesis; seeking effective therapeutics covering both liver and intestinal disease in PSC is of high clinical relevance.
- Independent of anti-cholestatic effects, NorUDCA has recently been shown to possess direct immunomodulatory properties on CD8^+^ T cell metabolism, lymphoblastogenesis and clonal expansion through targeting mTORC1 signaling.
- Since mTORC1 serves as critical metabolic checkpoint orchestrating T_H_17/Treg axis, inhibiting mTORC1 activity represents a potential treatment avenue counteracting T_H_17/Treg imbalance under intestinal inflammatory conditions.

*What are the new findings?:*

- NorUDCA enriches FOXP3^+^ Treg population in both liver and intestinal tissue in the cholestatic *Mdr2*^*–/–*^ mouse model of PSC.
- NorUDCA exhibits direct immunomodulatory efficacies in suppressing excess T_H_17 cell-mediated intestinal immunopathology and promotes FOXP3^+^ Treg generation in an experimental IBD model.
- Mechanistically, NorUDCA counteracts T_H_17/Treg imbalance by restricting glutaminolysis in differentiating T_H_17 cells, thus suppresses α-Ketoglutarate-dependent mTORC1 activation, glycolysis and enhances FOXP3 expression.
- NorUDCA’s impact on mTORC1 signaling was further confirmed in circulating CD4^+^ T cells from patients with PSC and IBD.

*How might it impact on clinical practice in the foreseeable future?:* These findings advance our current understanding of therapeutic potentials of NorUDCA, which might represent a novel therapeutic strategy in the treatment of PSC and concomitant IBD and other T_H_17-mediated intestinal diseases.

## Introduction

T helper cells 17 (T_H_17) are a subset of CD4^+^ T cells whose development is driven by expression of the signature transcription factor Retinoic acid orphan receptor-gamma (RORγt) when differentiated from naive CD4^+^ T cells and are known to contribute to inflammatory immune responses [1]. In contrast, regulatory T cells (Treg), expressing the Forkhead box P3 (FOXP3), have suppressive function and can dampen immune response [1]. The differentiation of T_H_17 and Treg cells is known to be reciprocally regulated, influenced by a direct antagonistic interaction between the master transcription factors, RORγt and FOXP3, and environmental cues [2]. The balance between T_H_17 cell differentiation and Treg generation shapes the outcome of tissue homeostasis and inflammation [1]. Under homeostatic condition, T_H_17 cells develop normally in the intestine and play a pivotal role in host defense against microbial pathogens, while IL17 produced by T_H_17 as well as by other cells such as, γδT cells [3] crucially maintains intestinal mucosal barrier integrity [4]. However, when T_H_17 immune responses become excessive, the balance between T_H_17 and Treg are compromised which can drive immunopathogenesis of several inflammatory and autoimmune disorders [5], including inflammatory bowel disease (IBD) [6, 7].

24-Nor-Ursodeoxycholic acid (NorUDCA) (recently renamed as Norucholic acid [8]) is a novel therapeutic modified bile acid which has demonstrated promising results in phase II clinical trials for treating primary sclersoing cholangitis (PSC) [9], a potentially fatal autoimmune disease of the liver and bile ducts so far lacking effective medical treatment [10]. NorUDCA is currently under evaluation for its long-term effect on PSC in a phase III trial (NCT03872921). Recently we have uncovered a yet unrecognized immunoregulatory role of NorUDCA in directly modulating CD8^+^ T cell metabolism and immune response through blunting mTORC1 activity during hepatic immunopathology, profoundly distinct from its parent compound UDCA [11]. This offers exciting entry points of exploring NorUDCA’s additional therapeutic potentials in applications of T cell mediated diseases, in addition to liver disorders. Since PSC is highly associated with IBD driven by T_H_17/Treg imbalance linked to dysregulated mTORC1 activity [12, 13, 14], we hypothesize that NorUDCA might harness therapeutic potentials in modulating T_H_17/Treg axis in intestinal inflammation.

We first assessed whether NorUDCA affects T_H_17/Treg tissue distribution in the cholestatic *Mdr2*^*–/–*^ mouse model of PSC. We further specifically investigated NorUDCA’s effect on modulating T_H_17/Treg balance in a CD4^+^ T cell driven IBD model induced by adoptive transfer of CD25^−^CD44^low^CD45RB^high^CD4^+^ T cells (CD4^+^ T_Naïve_) into *Rag2*^*–/–*^ mice. Further mechanistical studies were performed using molecular approaches, flow cytometry and metabolic assays in murine T_H_17 cells *in vitro*. Human peripheral CD4^+^ T cells from healthy volunteers and PSC patients with associated IBD were used to validate NorUDCA signaling effects observed in murine system.

## Methods

### *Mdr2(Abcb4)* ^*–/–*^ mouse model

Multidrug resistance gene 2 *Mdr2(Abcb4)* ^*–/–*^ (FVB/N background) male mice were fed±NorUDCA (0.5%, wt/wt) containing diets for 12 days starting at the age of 8 weeks after birth, a time point when sclerosing cholangitis is already fully established [15]. Wild-type (WT) FVB/N male mice received standard chow diet. Experiments were approved by the Austrian Federal Ministry for Science and Research at Medical University of Vienna (MUV) (BMWFW-66.009/0008-WF/V/3b/2015).

### Adoptive CD4^+^ T cell transfer colitis model

To induce colitis, CD4^+^ T_Naïve_ cells were sorted from C57BL/6N mice by FACS and 0.5×10^6^ cells were adoptively transferred into each *Rag2*^*–/–*^ recipient mouse which either receive standard chow or 0.5% (wt/wt) NorUDCA-supplemented diet upon transfer. At week6, mononuclear cells from tissues of T cell reconstituted *Rag2*^*–/–*^ mice were isolated, single cell suspensions were prepared as described in the supplementary material.

### Isolation of fecal bacterial microbiota and 16S rRNA gene sequencing analysis

DNA of the mouse fecal microbiota was isolated by using QIAamp Fast DNA Stool Mini Kit (Qiagen) according to the manufacturer’s instructions. 16S rRNA gene amplicon sequencing analysis was used to determine the alteration in gut microbiota composition as described previously [16].

### *In vitro* T_H_17 cell differentiation assay

Primary mononuclear cells were isolated from peripheral lymph nodes and spleens of C57BL/6N male mice (bred in the mouse facility of MUV) and magnetically enriched by for CD4^+^T_Naïve_ cells using Miltenyi Naïve CD4^+^ T cell isolation kit and submitted to T_H_17 culture as described in the supplementary material.

### Flow cytometric analysis

Cell surface and intracellular FACS stains for *ex vivo* isolated immune cells were performed at 4°C for 30min as previously described [11] using antibodies listed in Supplementary table 1. Intracellular staining of cytokines and transcription factors of *in vitro* cultured T_H_17 cells was performed as described in [11]. Intracellular staining of pRPS6^Ser235/236^ of *in vitro* cultured T_H_17 cells was performed by fixing for 10 min at 37°C with Cytofix (BD), permeabilized 20min on ice with methanol and stained with the indicated phospho-Abs for 30min in the dark at 4°C in PBS/2% FCS. For extracellular staining, T_H_17 cells were incubated with Abs for 30 min on ice.

### Glucose uptake assay

Murine T_H_17 cells differentiated in the presence or absence of NorUDCA were washed with PBS prior to incubation with 10μM D-glucose analog 2-[N-(7-nitrobenz-2-oxa-1,3-diazol-4-yl) amino]-2-deoxy-D-glucose (2-NBDG), a fluorescent analogue of glucose, for 30min at 37°C. Next cells were washed and submitted to flow cytometry for detection of fluorescence incorporated by the cells.

### Histopathology, PAS and multicolor-Immunofluorescence staining

Histopathology, Periodic Acid Schiff (PAS) staining for mucin and multicolor-Immunofluorescence Staining was performed as described previously [15].

### Serum analysis

Serum biochemistry was performed as described previously[17].

### Compound treatments

Concentrations of NorUDCA (500μM) for *in vitro* experiments were selected to mimic the *in vivo* serum bile acid level of mice on 0.5% (wt/wt) NorUDCA supplemented diet [11] (also as indicated in figure 2B). Concentrations of NorUDCA selected for *in vitro* experiments were previously tested not affecting viability in T cells [11]. Rapamycin (mTOR inhibitor; Cell signaling) was used as 100nM. Cell permeable *α*-KG (Sigma) was used as 1.5mM.

### Gene expression analysis

RNA isolation from primary murine T_H_17 culture, cDNA synthesis and real-time PCR were performed as described previously [17]. Gene expression was calculated relative to *Hprt*. Primer sequences were used as following: *Gls2* forward, 5’-AGCGTATCCCTATCCACAAGTTCA-3’; *Gls2* reverse, 5’-GCAGTCCAGTGGCCTTCAGAG-3’; *Hprt* forward, 5’-CTGGTGAAAAGGACCTCTCG-3’; *Hprt* reverse, 5’-TGAAGTACTCATTATAGTCAAGGGCA-3’.

### Human T cell experiments

3 independent experiments were performed with peripheral T cells obtained from 3 PSC patients with co-existing IBD (Mayo clinical score 0-2) and 3 age and gender matched healthy volunteers were performed following the Declaration of Helsinki and approved by the Ethics Committee of the MUV: 747/2011 and 2001/2018. 2 patients were newly diagnosed and only 1 patient was on UDCA treatment at the time of blood sampling. Bulk T cells from peripheral blood mononuclear cells (PBMCs) were isolated by negative depletion [18], labeled with 1μM CFSE and rested overnight in RPMI 1640 medium with 5% heat-inactivated FCS, 2mM L-glutamine, 100μg/ml streptomycin and 100U/ml penicillin. T cells were stimulated with plate-bound anti-CD3 (1μg/ml) plus soluble anti-CD28 (0.5μg/ml) Abs ± NorUDCA or Rapamycin. Lymphoblastogenesis, proliferation and mTORC1 activity were assessed on day 3 on a Fortessa flow cytometer (BD Biosciences) and quantified according to the CFSE peaks with FlowJo (BD) [18]. Clones/antibodies are shown in Supplementary Table 1.

### Quantification and statistical analysis

All values are expressed as mean ± standard error and were statistically analyzed as detailed in the figure legends using GraphPad Prism v.7.0 (La Jolla, CA, USA). A *P* value of less than 0.05 was considered statistically significant and indicated as follows: *= *P*<0.05, **= *P*<0.01, ***= *P*<0.001, ****=*P*<0.0001.

## Results

### NorUDCA preferentially enriches FOXP3^+^ Tregs in both liver and intestinal tissue in the cholestatic *Mdr2*^−/–^ mouse model of sclerosing cholangitis

Previously we have reported that 4-week NorUDCA feeding reversed sclerosing cholangiopathy in *Mdr2*^−/–^ mice by inducing bicarbonate-rich bile flow which reduces bile acid toxicity and reinforces the biliary “bicarbonate umbrella” [17, 19]. In a recent study, we have shown that short-term (12 days) NorUDCA feeding on 8-week-old *Mdr2*^−/–^ mice with fully established hepatobiliary injury and fibrosis was sufficient to demonstrate pronounced immunomodulatory efficacies reducing hepatic infiltration of innate and adaptive immune cells including both CD8^+^ and CD4^+^ T cells [11]. However, the potential impact of NorUDCA on T_H_17/Treg tissue distribution in this cholestatic *Mdr2*^−/–^ model has so far not been explored. Therefore, we analyzed how NorUDCA affects T_H_17/Treg tissue distribution in the *Mdr2*^−/–^ model by assessing the expression of signature transcription factor RORγt and FOXP3. Hepatic and intestinal CD4^+^ T cells isolated from *Mdr2*^−/–^ mice only showed minimal expression of RORγt in comparison to control wild-type mice, which was not altered by NorUDCA (data not shown). Interestingly, both hepatic and small intestinal intraepithelial FOXP3^+^ Tregs were expanded by NorUDCA in *Mdr2*^−/–^ mice (figure 1A-C) indicating that NorUDCA might have therapeutic potentials in promoting Treg generation *in vivo*.

**Figure 1.**
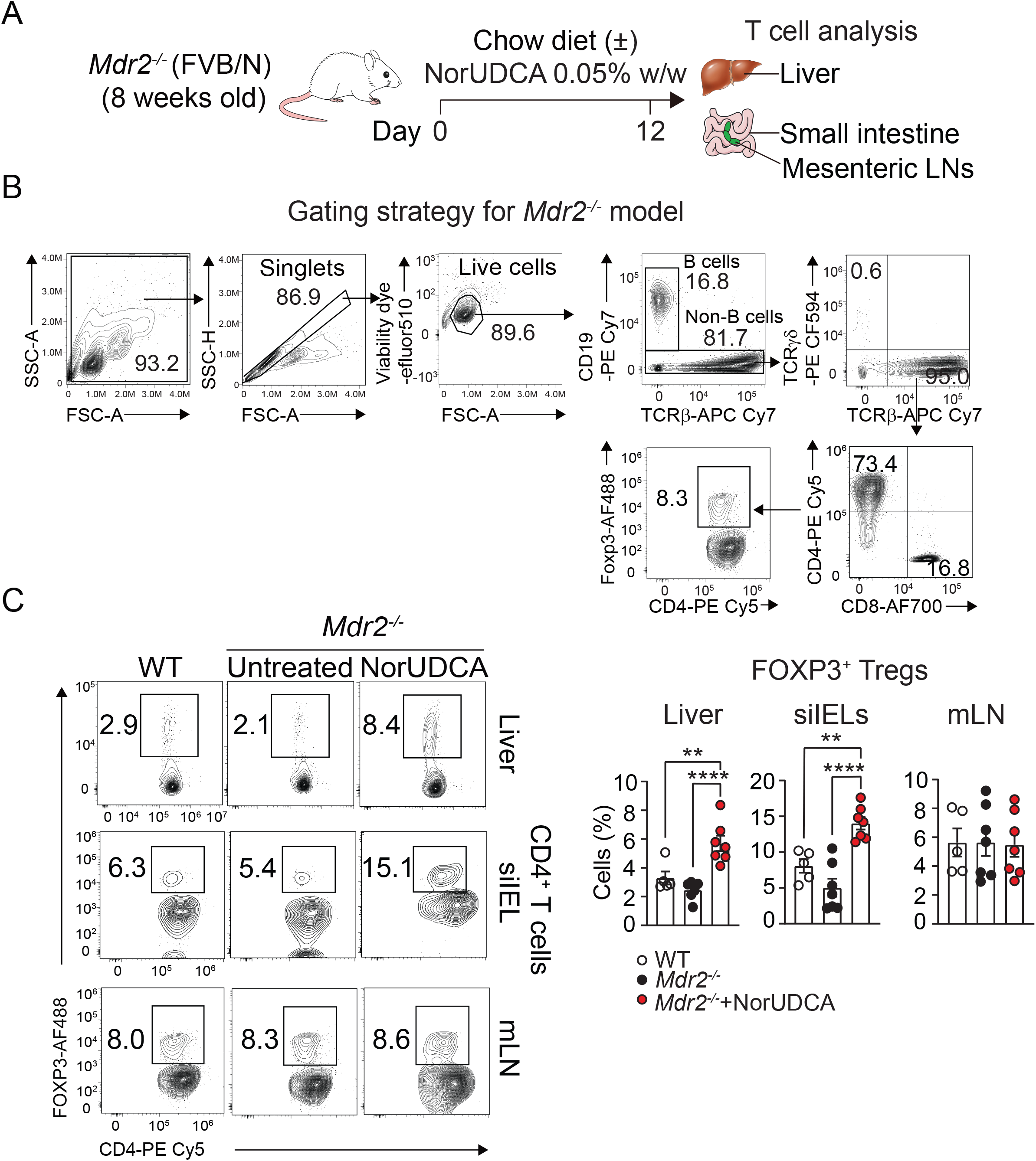
Short-term NorUDCA treatment enriches Tregs in both liver and intestine in cholestatic *Mdr2*^*–/–*^ model of spontaneous sclerosing cholangiopathy. (A) Experimental scheme. (B) Gating strategy for the analysis of *ex vivo* isolated T cells in *Mdr2*^*-/-*^ model. (C) Representative and quantitative analysis of Treg distribution within indicated tissues. Data are representative of 3 independent experiments. At least 5 animals were used per group during experiments. Quantitative data are presented as mean±SE. *P* values were calculated by one-way ANOVA corrected for multiple comparisons with Dunnett test. **= *p*<0.01, ****= *p*<0.0001. siIEL, small intestinal intraepithelial lymphocytes; mLN, mesenteric lymph nodes.

### NorUDCA alleviates intestinal immunopathology and modulates gut microbiome in adoptive CD4^+^ T cell transfer colitis model of IBD

The extent of the gut inflammatory response in the cholestatic *Mdr2*^−/–^ mouse model of PSC is dependent on the microbiota composition which varies among different animal facilities [20]. Since *Mdr2*^−/–^ mice housed at our animal facility lacked overt intestinal inflammation reported in other studies [21], we specifically investigated NorUDCA’s effect on modulating T_H_17/Treg balance in a well-established CD4^+^ T cell driven IBD model induced by adoptive transfer of CD4^+^ T_Naïve_ cells into recipient *Rag2*^−/–^ mice lacking T and B cells. Following interaction with intestinal antigen presenting cells, the transferred CD4^+^ T_Naïve_ cells mainly differentiate into pathogenic T_H_1 and T_H_17 cells which express IFN*γ* [22] causing subsequent progressive intestinal tissue infiltration and inflammation mimicking immunopathological features of human IBD [23]. Importantly, the compromised T_H_17/Treg balance observed in the IBD patients [5, 24] is also seen in this adoptive CD4^+^ T cell transfer colitis model [25]. To investigate the impact of NorUDCA on modulating T_H_17/Treg balance *in vivo*, we performed NorUDCA treatment in this disease model during the entire course of the colitis development (figure 2A). Increased plasma bile acid levels indicated NorUDCA absorption and systemic enrichment *in vivo* (figure 2B). Compared with untreated mice, NorUDCA treatment significantly protected colitis mice against shortening of colon length (figure 2C), while suppressed immune cell infiltration (figure 2D), goblet cell loss (figure 2E) and preserved intestinal architecture and mucus barrier integrity as manifested by tissue staining (figure 2D-F).

**Figure 2.**
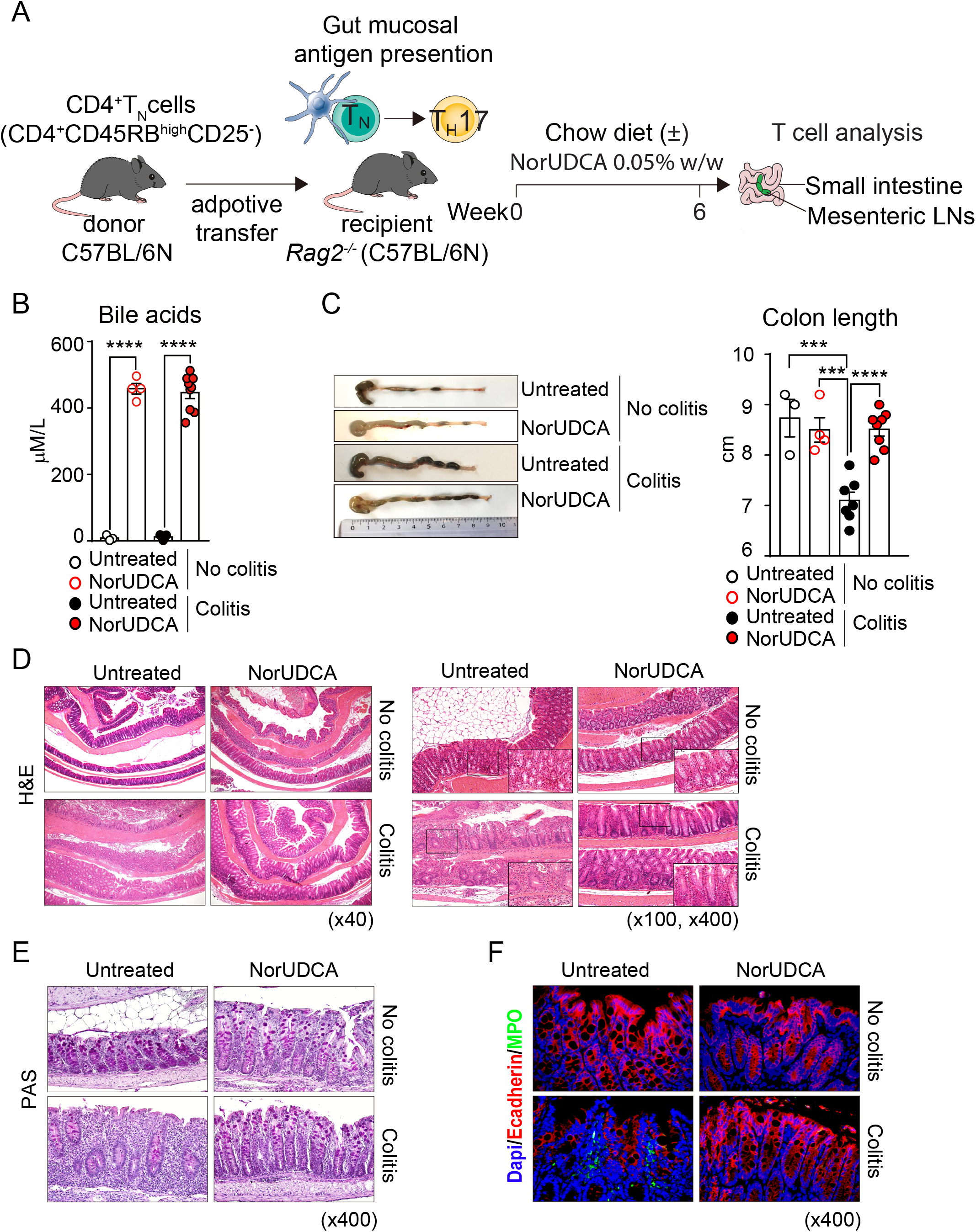
NorUDCA alleviates intestinal immunopathology in a CD4^+^ T cell driven colitis model of IBD. (A) Experimental scheme. (B) Serum bile acids. (C) Representative and quantitative analysis of colon length of indicated groups. (D) Representative H&E staining of colon sections are shown. (E) Representative PAS staining for mucin of colonic goblet cells. (F) Representative multicolor-immunofluorescence staining of Ecadherin and MPO of colonic slides of indicated groups. Data are summary of 2 independent experiments. At least 4 animals were used per group with colitis. Quantitative data are presented as mean±SE. *P* values were calculated by one-way ANOVA corrected for multiple comparisons with Dunnett test. ***= *p*<0.001, ****= *p*<0.0001. H&E, hematoxylin and eosin; PAS, periodic acid schiff; MPO, myeloperoxidase.

Gut microbiome and dysbiosis play important roles in the development of IBDs [26]. Therefore, we next examined whether NorUDCA treatment modulated the composition of gut microbiome in colitis mice. Analysis of fecal samples by 16S ribosomal RNA gene sequencing showed that NorUDCA treatment significantly restored bacterial diversity in colitis mice as reflected by the increase of Shannon index and relative abundance of *Akkermansia muciniphila* (known to be linked with protective intestinal barrier functions [27]) and *Lactobacillus* (known for beneficial roles in both IBD animal models and patients [28]) which are key species of human intestinal bacteria prominently decreased in patients with IBDs (see supplementary figure 1). Together, these data indicate pleiotropic beneficial mode of actions of NorUDCA in counteracting inflammatory responses and intestinal dysbiosis during intestinal inflammation.

### NorUDCA suppresses T_H_17 cell intestinal infiltration and promotes Treg expansion in adoptive CD4^+^ T cell transfer colitis model of IBD

We next evaluated the therapeutic efficacy of NorUDCA on T_H_17/Treg tissue infiltration and T_H_17 cell effector functions by assessing expression of key proinflammatory cytokines such as IL-17A, IFNγ and IL-22 [25] (figure 3A for gating strategy). Flow cytometry analysis revealed that the frequency of RORγt^−^FOXP3^+^ Treg population among intraepithelial lymphocytes, lamina propria lymphocytes and mesenteric lymphocytes were increased in the colitis mice treated with NorUDCA (figure 3B and C). This is accompanied with a reduction of frequencies of intraepithelial- and lamina propria-infiltrating CD4^+^ T cells double producing IL-17A/IFNγ and IL-17A/IL-22 in the colitis mice treated with NorUDCA (figure 3B and C). Taken together, our results suggest that NorUDCA treatment counteracted the development of T_H_17/Treg imbalance in the setting of intestinal inflammation.

**Figure 3.**
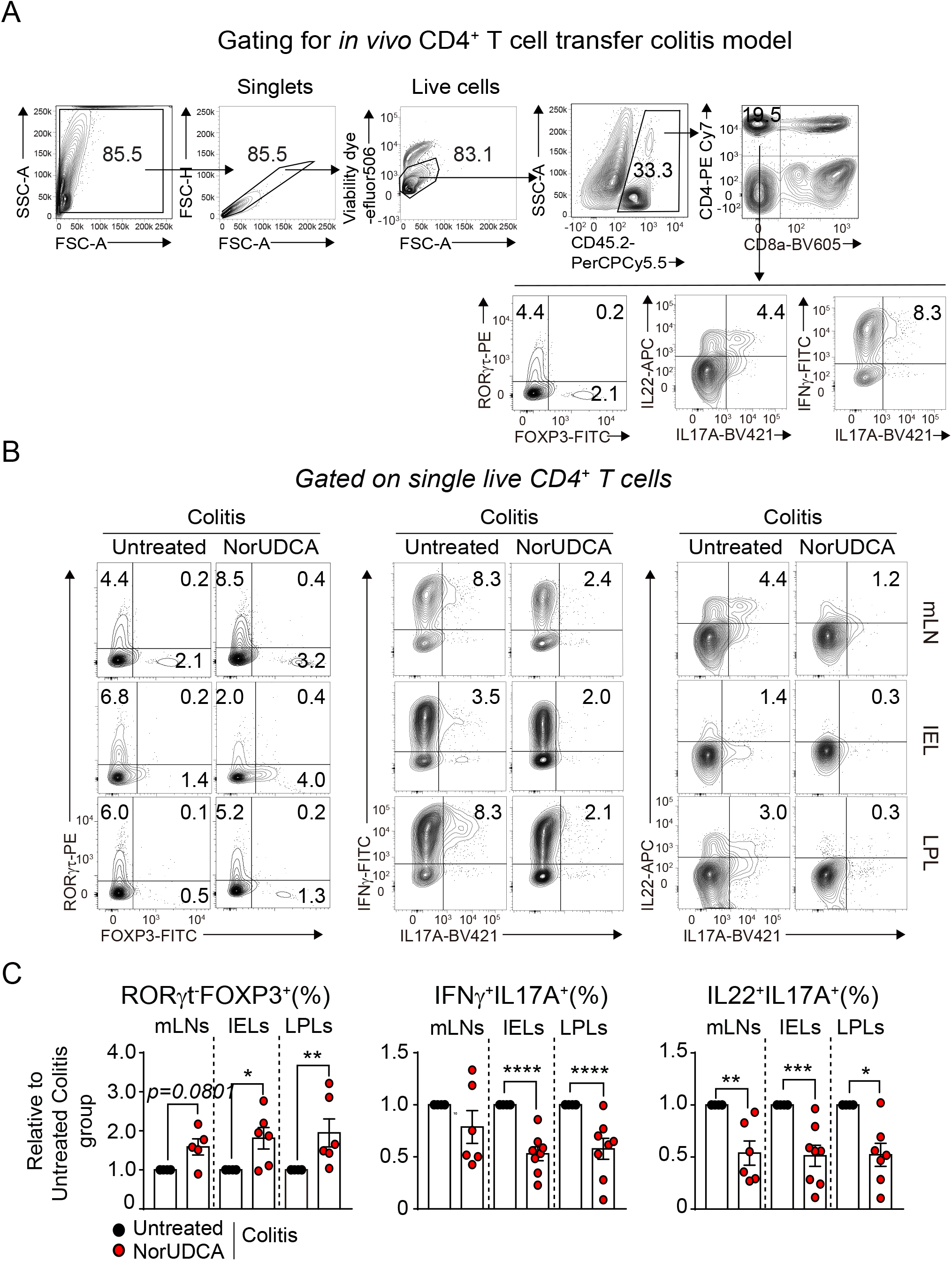
NorUDCA suppresses T_H_17 cell intestinal infiltration and promotes Treg expansion in the adoptive CD4^+^ T cell transfer colitis model of IBD. (A) Gating strategy for the analysis of *ex vivo* isolated T cells in CD4^+^ T cell adoptive transfer colitis model. (B) Representative plot of infiltrating CD4^+^ T cells expressing indicated transcription factors and cytokines of T_H_17 cells in indicated tissues. (C) Quantitative analysis of (B). Data are summary of 2 independent experiments. At least 3 animals were used per group with colitis. Quantitative data are presented as mean±SE. *P* values were calculated by one-way ANOVA corrected for multiple comparisons with Dunnett test. ***= *p*<0.001, ****= *p*<0.0001. mLN, mesenteric lymph nodes; siIEL, small intestinal intraepithelial lymphocytes; LPL, lamina propria lymphocytes.

### NorUDCA represses T_H_17 cell differentiation and is associated with alterations in mTORC1 activity and decreased glycolysis

The immunomodulatory efficacy of NorUDCA on T_H_17/Treg axis *in vivo* incited us to thoroughly study whether NorUDCA directly affects the differentiation potential of T_H_17 cells. Therefore, we employed a T_H_17 *in vitro* system by activating and differentiating murine primary CD4^+^ T_Naïve_ cells towards T_H_17 cells in the presence or absence of NorUDCA. T_H_17 cell differentiated in the presence of NorUDCA exhibited normal RORγt expression, however, less IL-17A expression on RORγt^+^ cells and lower frequency of cells double producing the signature cytokines IL-17A and IL-17F, indicating NorUDCA repressed T_H_17 cell effector function (figure 4A). Intriguingly, we detected an increased frequency of FOXP3 expressing cells under T_H_17 differentiation condition in presence of NorUDCA compared to the untreated control cells (figure 4A), suggesting a conversion of T_H_17 cells into Tregs.

**Figure 4.**
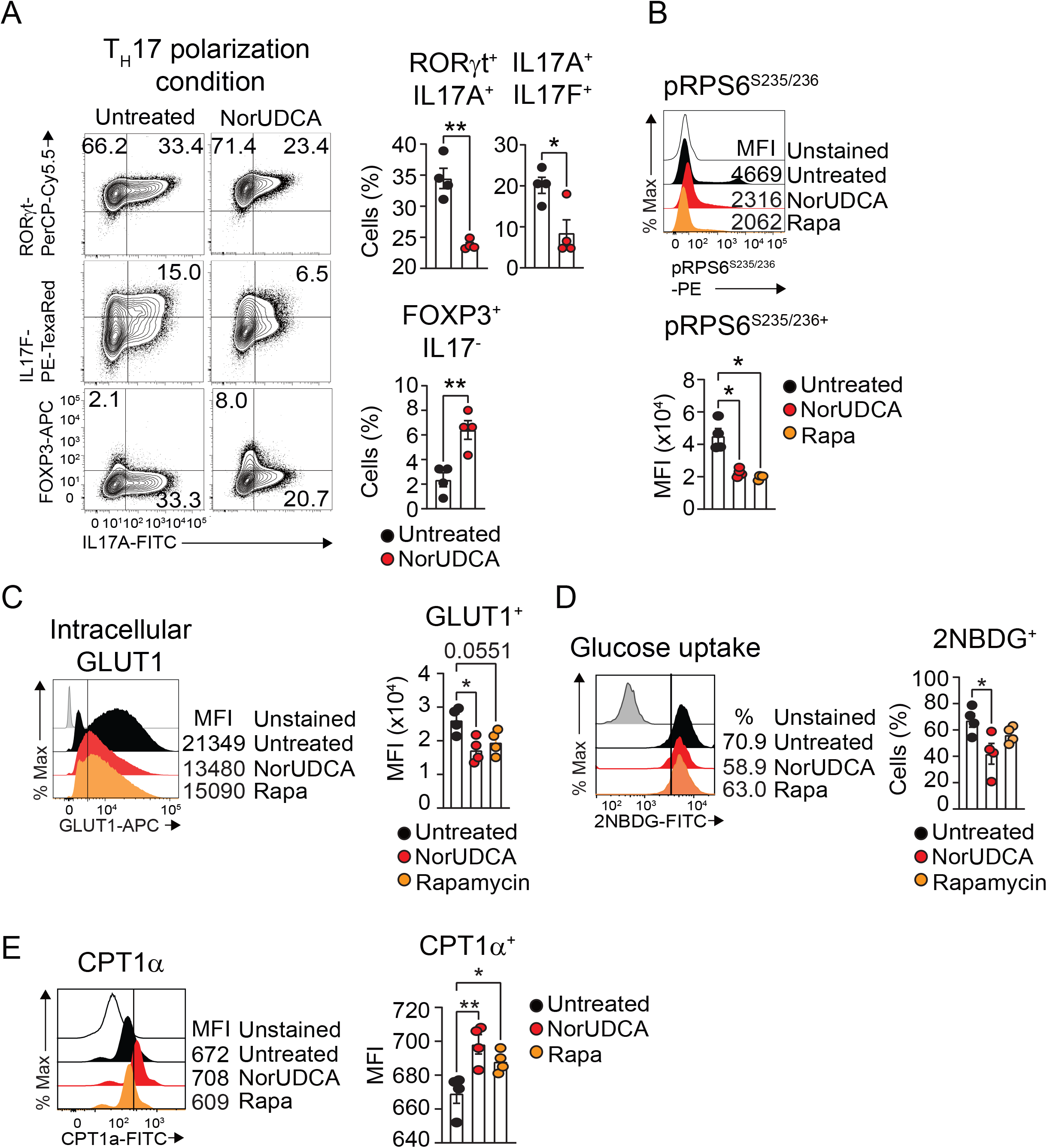
NorUDCA represses T_H_17 cell differentiation and is associated with alterations in mTORC1 activity and decreased glycolysis. (A) Representative contour plots and quantitative analysis of *in vitro* differentiated T_H_17 cells expressing signature transcription factors and cytokines. (B) Representative histograms and quantitative analysis of intracellular pRPS6^Ser235/236^ expression on differentiating T_H_17 cells. (C) Representative histograms and quantitative analysis of total (surface and intracellular) GLUT1 expression on differentiating T_H_17 cells. (D) Glucose uptake analyzed on differentiating T_H_17 cells. (E) Representative histograms and quantitative analysis of intracellular CPT1α expression on differentiating T_H_17 cells. Data are summary of 3 independent experiments. Quantitative data are presented as mean±SE. *P* values were calculated by one-way ANOVA corrected for multiple comparisons with Dunnett test using untreated group as reference. ***= *p*<0.001, ****= *p*<0.0001. Rapa, Rapamycin.

The mammalian target of rapamycin (mTOR) is an evolutionarily conserved serine/threonine kinase that senses and integrates diverse immune signals and metabolic cues for activation and lineage commitment in T cells [29]. mTOR is part of two distinct multiprotein complexes, mTORC1 and mTORC2, which are modulated by several upstream signaling pathways and exerts different functions [30]. mTORC1 serves as a central regulator mediating lineage decisions between T_H_17 cells and Tregs. The inhibition of mTORC1 activity blocks the generation of T_H_17 cells [13], while activation of mTORC1 promotes the generation of FOXP3^+^ Tregs [31]. Thus, we investigated whether NorUDCA impacts on mTORC1 activity during T_H_17 cell differentiation. Kinase complex mTORC1 licenses initiation and elongation of translation via ribosomal protein S6 kinase 1 (S6K1) which phosphorylates Ribosomal protein S6 (RPS6) [30]. As a downstream readout of mTORC1 activity, we evaluated the intracellular levels of phosphorylation of RPS6 at Ser235/236 (pRPS6^Ser235/236^) by flow cytometry. Rapamycin was included as mTORC1 inhibition control. NorUDCA decreased the expression of pRPS6^Ser235/236^ among T_H_17 cells to the level of Rapamycin (figure 4B) indicating that mTORC1 activity was inhibited by NorUDCA during T_H_17 cell differentiation.

Given that mTORC1 acts as a critical bioenergetic checkpoint orchestrating cellular metabolisms in T_H_17 cells during differentiation [29], we next investigated if NorUDCA induced mTORC1 inhibition has an impact on metabolic pathways of T_H_17 cells. Since glycolysis in T cells is positively controlled by mTORC1, we next examined if NorUDCA alters glycolytic activity in T_H_17 cells. As a consequence of mTORC1 inhibition, the expression of glucose transporter 1 (GLUT1) (key transporter responsible for glucose uptake in T cells [32]) was repressed by NorUDCA in differentiating T_H_17 cells (figure 4C). A glucose uptake assay revealed that differentiating T_H_17 cells treated with NorUDCA incorporated less glucose analogue compared to untreated cells, suggesting a downregulation of glycolysis by NorUDCA (figure 4D). Conversely, expression of carnitine palmitoyltransferase 1α (CPT1α), a rate limiting enzyme in fatty acid oxidation (FAO) [33], was upregulated by NorUDCA in T_H_17 cells (figure 4E), supporting the evidence that inhibition of glycolysis enforces fatty acid oxidation in CD4^+^ T cells [34].

### NorUDCA restricts glutamine metabolism that licenses mTORC1 activation in differentiating T_H_17 cells

Integrating with glycolysis, T_H_17 cells accelerate glutamine uptake and glutaminolysis to meet the bioenergetic demands mounting for rapid expansion and immune response [35]. Glutamine is a conditionally essential amino acid consumed in fast proliferating cells [36]. Activated T cells upregulate amino acid transporters and enzymes that metabolize glutamine [37]. Glutamine is initially hydrolyzed via the enzyme glutaminase (GLS) to produce glutamate which is used in protein synthesis and generation of glutathione to regulate reactive oxygen species and exchanged to promote cystine uptake [37]. Glutamate is further metabolized to a-ketoglutarate, which provides anaplerotic support of mitochondrial tricarboxylic acid cycle [37]. Glutamine deprivation was shown to impair T_H_17 development and promote FOXP3 expression in a process that is linked to mTORC1 signaling [35]. Therefore, we investigated if NorUDCA has an impact on glutamine metabolism / glutaminolysis in T_H_17 cells. We observed that surface expression of major glutamine transporter ASCT2 (also known as Slc1A5) [38] was not affected by NorUDCA in *in vitro* generated T_H_17 cells (figure 5A). However, the gene expression level of *Gls2*, the rate-limiting enzyme participating in glutaminolysis [39], was strongly downregulated in NorUDCA treated T_H_17 cells which coincided with lower mTORC1 activity (figure 5B and C). Interestingly, supplement of cell permeable α-ketoglutarate abrogated mTORC1 inhibition induced by NorUDCA in differentiating T_H_17 cells observed under glutamine deprivation condition, which also held true for NorUDCA’s suppression on enhancement of FOXP3 expression but not IL17A expression (figure 5C and D). Moreover, we found that the ability of NorUDCA to reduce GLUT1 expression is also positively correlated with its modulation on glutaminolysis activity as evidenced that GLUT1 expression is restored by adding α-ketoglutarate back to the glutamine deprivation condition in the presence of NorUDCA (figure 5E). Taken together, our finding suggests that NorUDCA alters metabolic reprogramming that conditions the balance between T_H_17 and FOXP3^+^ Tregs.

**Figure 5.**
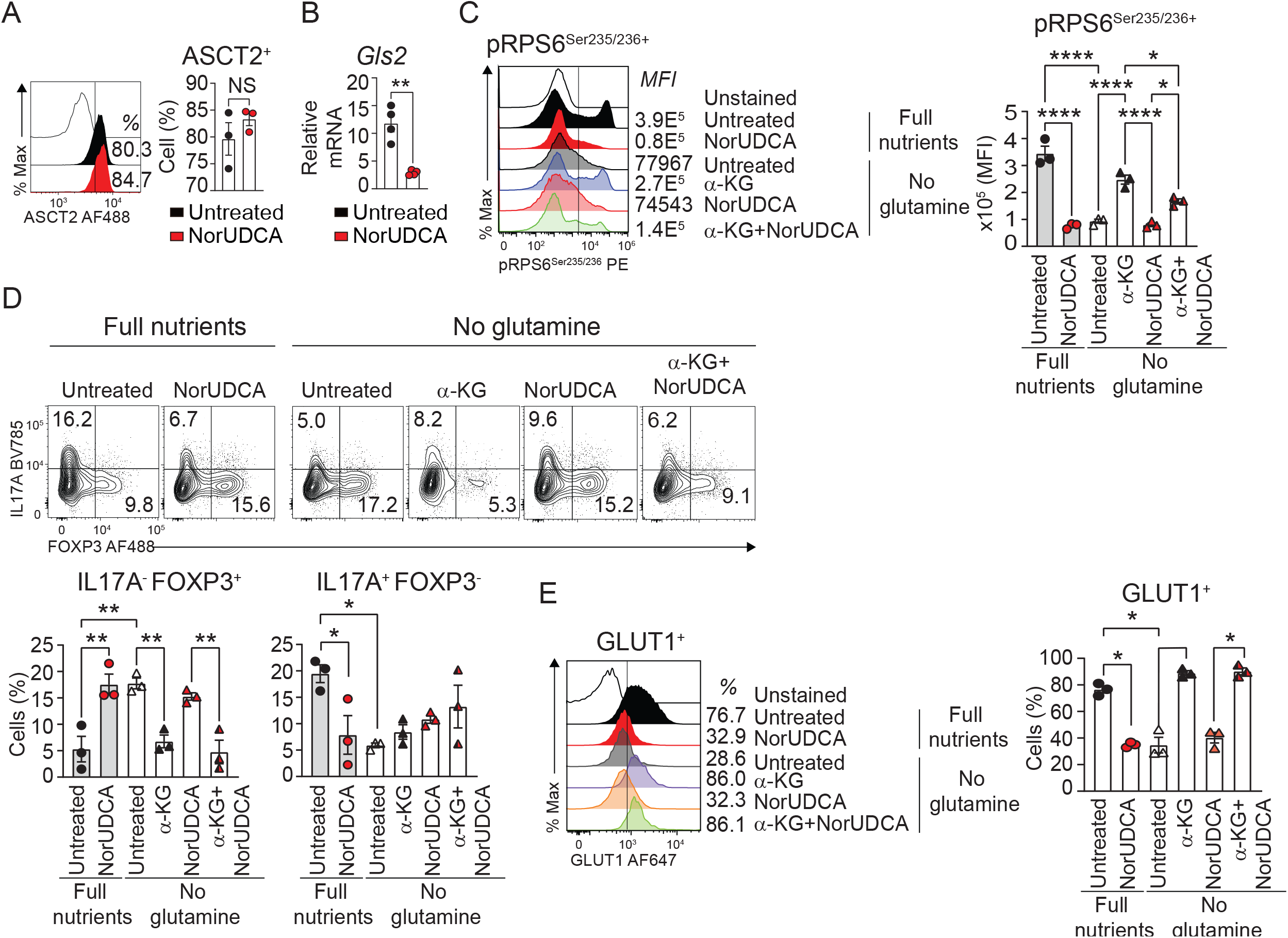
NorUDCA restricts glutamine metabolism that licenses mTORC1 activation in differentiating T_H_17 cells. (A) Representative histograms and quantitative analysis of ASCT2 expression on T_H_17 cells. (B) Quantitative RT-qPCR of expression of *Gls2* (normalized to house-keeping *Hprt*). (C) Representative histograms and quantitative analysis of intracellular pRPS6^Ser235/236^ expression on differentiating T_H_17 cells of indicated conditions. (D) Representative contour plots and quantitative analysis of T_H_17 cells expressing indicated transcription factors and cytokines. (E) Representative histograms and quantitative analysis of total (surface and intracellular) GLUT1 expression on differentiating T_H_17 cells of the indicated conditions. Data are summary of 3 independent experiments. Quantitative data are presented as mean±SE. *P* values were calculated by one-way ANOVA corrected for multiple comparisons with Dunnett test using untreated group as reference. **= *p*<0.01, ***= *p*<0.001, ****= *p*<0.0001. Rapa, Rapamycin; a-KG, a-ketoglutarate.

### NorUDCA limits robust lymphoblastogenesis and clonal expansion in PSC-IBD patient-derived circulating CD4^+^ T cells with hampered mTORC1 kinase activity

Finally, we investigated whether some of our key findings obtained with murine cells apply to human peripheral CD4^+^ T cells especially peripheral CD4^+^ T cells from patients with PSC and IBD (PSC-IBD), where dysregulated T cells are associated with disease progression. We examined the impact of NorUDCA on CD4^+^ T cell activation, blasting, clonal expansion and mTORC1 *in vitro* by *ex vivo* activation of isolated peripheral T cells from healthy volunteers and PSC-IBD patients in the absence or presence of NorUDCA. Rapamycin was used as control for mTORC1 inhibition. Compared with CD4^+^ T cells from healthy volunteers, CD4^+^ T cells from PSC-IBD patients demonstrated robust potentials of lymphoblastogenesis (figure 6A), clonal expansion (figure 6B) and mTORC1 activity (figure 6C). In accordance to our observations gained from the murine experiments, NorUDCA reduced cell size and clonal expansion in CD4^+^ T cells from PSC-IBD patients while not affecting cell viability and activation (see supplementary figure 2). We also investigated if NorUDCA affects mTORC1 activity in PSC-IBD CD4^+^ T cells by assessing expression of pRPS6. Following NorUDCA treatment, we detected decreased lymphoblastogenesis (figure 6D) and clonal expansion (figure 6E) in parallel with lower mTORC1 activity in proliferating CD4^+^ T cells of PSC-IBD patients. Taken together, these data confirmed that NorUDCA’s mTORC1 inhibitory effect observed in murine CD4^+^ T cells was also seen in the circulating CD4^+^ T cells from PSC-IBD patients.

**Figure 6.**
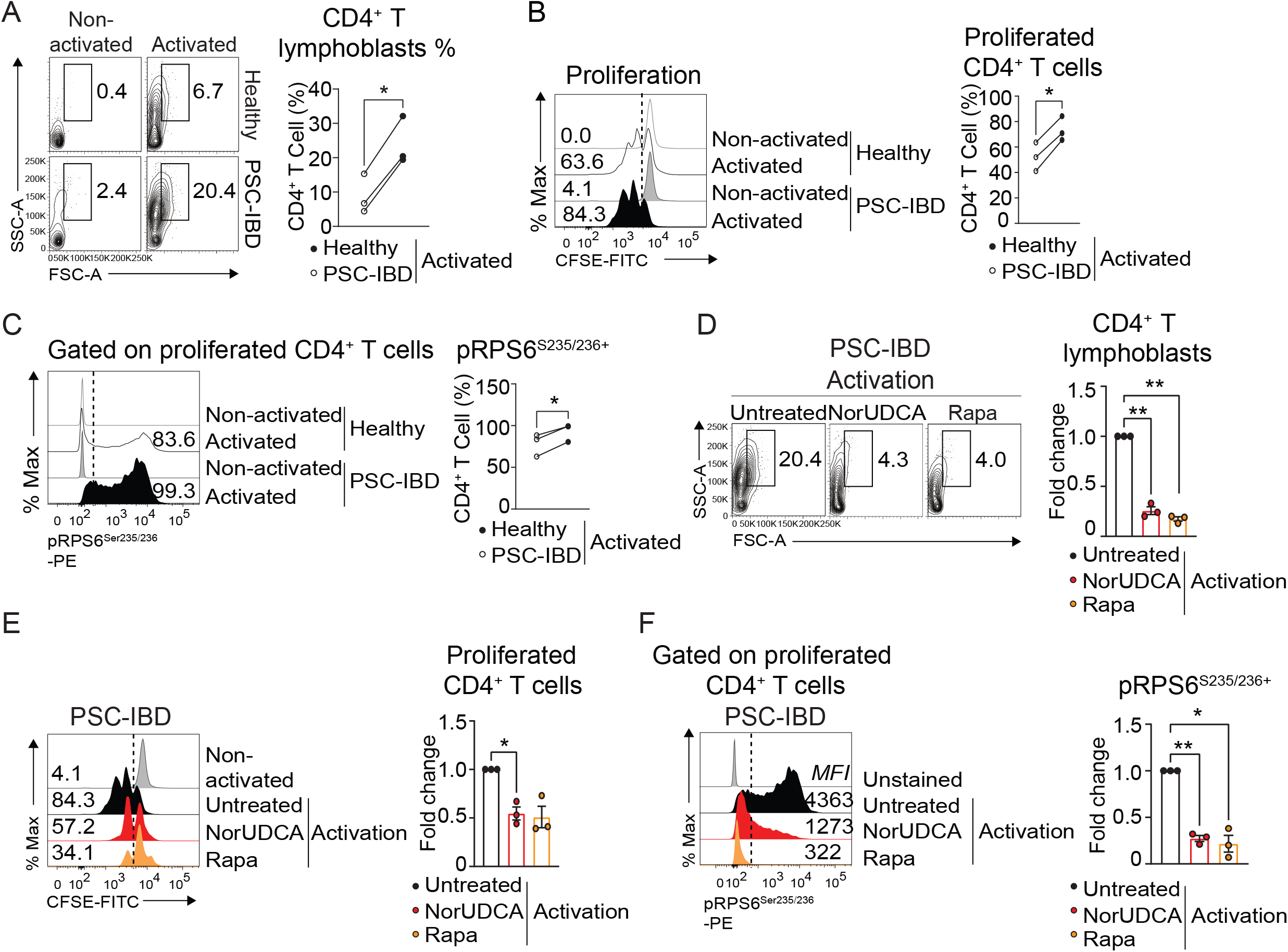
NorUDCA reduces lymphoblastogenesis and clonal expansion in circulating PSC-IBD CD4^+^ T cells with suppressed mTORC1 kinase activity. (A) Circulating human T cells from patients with PSC and associated IBD (PSC-IBD) as well as age- and gender-matched healthy volunteers were activated. (A, B) Representative plots showing lymphoblastogenesis and expansion of CD4^+^ T cells under indicated conditions. Quantitative analysis are shown in the right panel. (C) pRPS6^Ser235/236^ expression on proliferating CD4^+^ T cells from healthy volunteers and PSC-IBD patients are shown. (D-E) Effects of NorUDCA on lymphoblastogenesis (D) and expansion (E) of CD4^+^ T cells from PSC-IBD patients are shown. (F) pRPS6^Ser235/236^ expression on proliferating PSC-IBD CD4^+^ T cells under indicated conditions are shown. Quantitative analysis in (D, E, F) were performed using untreated group as reference and are shown in the right panel. Data are summary of 3 independent experiments. Quantitative data are presented as mean±SE. *P* values were calculated by one-way ANOVA corrected for multiple comparisons with Dunnett test using untreated group as reference. **= *p*<0.01, ***= *p*<0.001, ****= *p*<0.0001. PSC-IBD, primary sclerosing cholangitis concomitant with inflammatory bowel diseases; Rapa, Rapamycin.

## Discussion

PSC is an autoimmune cholestatic liver disease highly associated with IBD in about 70% and so far lacks effective medication [40]. Therefore, seeking therapeutics targeting both liver and intestinal disease for PSC and concomitant IBD is of high clinical interest. NorUDCA as novel therapeutic approach for treating PSC has been tested clinically with promising results in improving cholestasis [9]. In a recent study, we uncovered a direct anti-inflammatory and immunomodulatory role of NorUDCA in reshaping metabolic and (phospho)proteomic landscape in CD8^+^ T cells, independent of its well-established anti-cholestatic efficacy [11]. Notably, UDCA, the structurally closest related naturally occurring endogenous bile acids and parent compound of NorUDCA, entirely lacked the signaling effects and immunometabolic modulatory functions throughout our comprehensive study comparing NorUDCA and UDCA *in vitro* and *in vivo* [11]. These interesting findings incited us to investigate if NorUDCA’s immunomodulatory mode of action might also apply to other immune cells, such as CD4^+^ T cells, and whether NorUDCA possesses therapeutic potentials in treating concomitant IBD.

Here, we report that in the cholestatic *Mdr2*^*–/–*^ mouse model of PSC where NorUDCA has been initially tested for its therapeutic efficacy on hepatobiliary injury and fibrosis [17], NorUDCA preferentially enriched hepatic and intestinal Treg populations. We further specifically investigated NorUDCA’s effect on counteracting T_H_17/Treg imbalance which is linked to pathogenesis of IBD and PSC [6, 7, 41, 42] in a CD4^+^ T cell driven colitis model induced by adoptive transfer of CD4^+^ T_Naïve_ cells into *Rag2*^*–/–*^ mice which closely mimic human IBD [23]. In this setting, we uncovered that NorUDCA possesses a yet-unrecognized direct modulatory potency on counteracting T_H_17/Treg imbalance in the gut and mitigates intestinal immunopathology induced by an excessive T_H_17 proinflammatory immune response. Our study for the first time demonstrates that beyond its therapeutic efficacy in improving hepatic inflammation and biliary injury, NorUDCA is able to exert direct immunomodulatory and anti-inflammatory effects *in vivo* outside the liver.

One key mechanism we discovered is that NorUDCA has a direct impact on T_H_17 cell differentiation and metabolism. Given the complexity of immune regulation under conditions of intestinal inflammation, we certainly cannot rule out the role of additional beneficial mechanisms induced by NorUDCA, such as impact on T_H_17 tissue migration, modulation of innate immunity or the observed alterations in the gut microbiome composition in the CD4^+^ T cell driven colitis model, which definitely requires further independent studies. However, by focusing on T_H_17 cells we further identified that through restricting glutaminolysis NorUDCA inhibits mTORC1 activity which is directly controlled by α-Ketoglutarate, a key metabolic intermediate biosynthesized along the glutaminolysis cascade (figure 7). Revisiting the mass spectrometry data set of NorUDCA-treated CD8^+^ T cells published in our previous study [11], we found that in accordance with our observation in T_H_17 cells the abundance of glutaminolysis rate-limiting enzyme GLS2 (Glutaminase 2) was also reduced. This suggests that GLS2 might be a potential metabolic checkpoint for NorUDCA’s modulatory actions of glutamine metabolism in adaptive immune cells, thus provides an interesting direction for future investigations. mTORC1 inhibition provides a crucial immunometabolic link controlling T_H_17 cell effector function [43, 44], modulating glycolysis and FAO [45]. Following mTORC1 inhibition, NorUDCA induces metabolic remodeling in T_H_17 cells with decreased glycolysis and enhanced FAO machinery, which ultimately leads to repressed effector function (figure 7). The metabolic reprogramming induced by NorUDCA in differentiating T_H_17 cells is concomitant with lineage plasticity as manifested by an increasing population of cells expressing FOXP3 suggesting counteraction of T_H_17/Treg imbalance.

**Figure 7.**
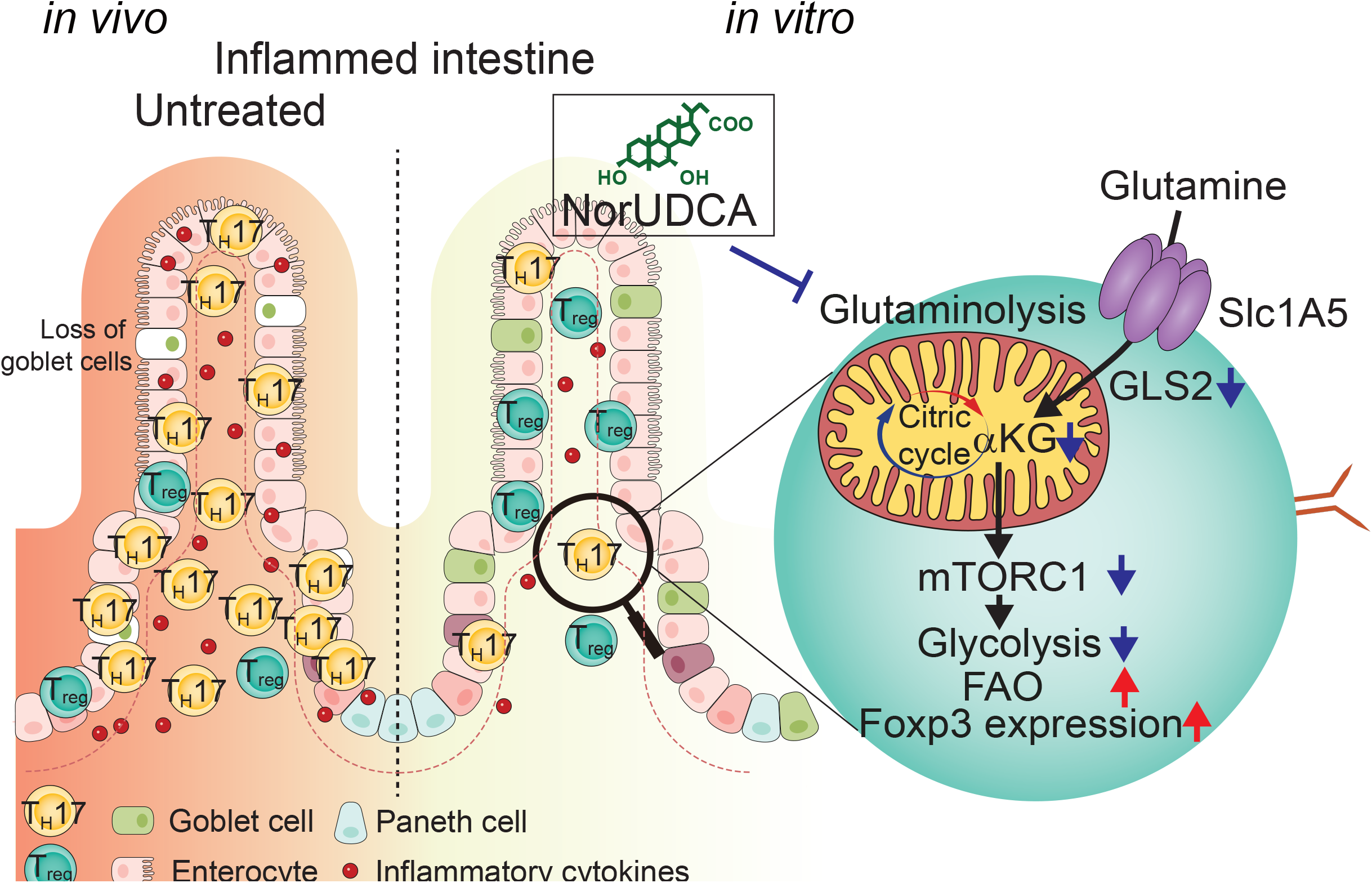
Graphical abstract. Schematic overview of how NorUDCA affects T_H_17/Treg balance *in vivo* and intracellular metabolism and signaling pathways in differentiating T_H_17 cells *in vitro*.

Our mechanistic studies also for the first time unveil that in addition to impacting on the classic molecular upstream transduction network regulating mTORC1, such as Ras-Erk-P90RSK previously published [11], NorUDCA can modulate metabolic upstream signaling pathways that licenses mTORC1 activation to exert subsequential alterations of metabolism and effector functions. Our original discovery about NorUDCA reshapes signal transduction cascades and metabolism at multiple dimensions may be a valuable addition to elucidate mechanism underlying NorUDCA’s immunomodulatory mode of actions. Thus, our current study provides a new conceptual framework of understanding metabolic regulatory properties of NorUDCA warranting future full-scale metabolic studies on NorUDCA in different T cell subsets, which should further advance our mechanistic understanding and potentially extend clinical applications of NorUDCA.

One of our key discoveries shows that circulating CD4^+^ T cells from PSC patients with IBD exhibit a hyperproliferative phenotype with enhanced mTORC1 activity, in contrast to CD4^+^ T cells from healthy volunteers. *In vitro*, NorUDCA suppressed mTORC1 in CD4^+^ T cells from PSC patients with IBD as efficiently as Rapamycin further validating our signaling findings made in the murine system. Interestingly, the mTORC1 hyperactivation status had also been detected in circulating CD8^+^ T cells in PSC patients in our recent published study [11]. Thus, this raises the questions whether mTORC1 hyperactivity seen in circulating T cells is also observed in tissue-infiltrating T cells in PSC or associated IBD. mTORC1 hyperactivation is implicated in the pathogenesis of several autoimmune diseases [46], however whether mTORC1 hyperactivity is linked with augmented pathogenicity perpetuating disease progression needs further investigation. Given that mTORC1 serves as critical metabolic checkpoint in immune cells and inflammation and metabolism are inextricably linked, future studies profiling the metabolic signatures of key immune cells driving pathogenesis of PSC or associated IBD may be justified. Many current medications for PSC or PSC with IBD lack specificity and are associated with significant off-target effects, highlighting the urgent need for exploring novel approaches. As such, NorUDCA by potentially targeting metabolic signaling pathways, might help to improve treatment efficacy for these disorders.

Taken together, here we demonstrate pronounced immunomodulatory efficacy of NorUDCA on counteracting T_H_17/Treg imbalance resulting in alleviated intestinal inflammation. Thus, our findings may have imminent therapeutic implications for treatment of PSC and associated IBD, where NorUDCA has shown promising results at least in part for the liver disease [9], while UDCA has no established efficacy [47]. Further studies are warranted to explore the array of potential therapeutic applications of NorUDCA in T_H_17-mediated intestinal and hepatic diseases.

