## Supplementary figures and images for "24-Nor-ursodeoxycholic acid counteracts T_H_17/Treg imbalance and ameliorates intestinal inflammation by restricting glutaminolysis in differentiating T_H_17 cells"

### Supplementary figures

#### Supplementary Fig. 1

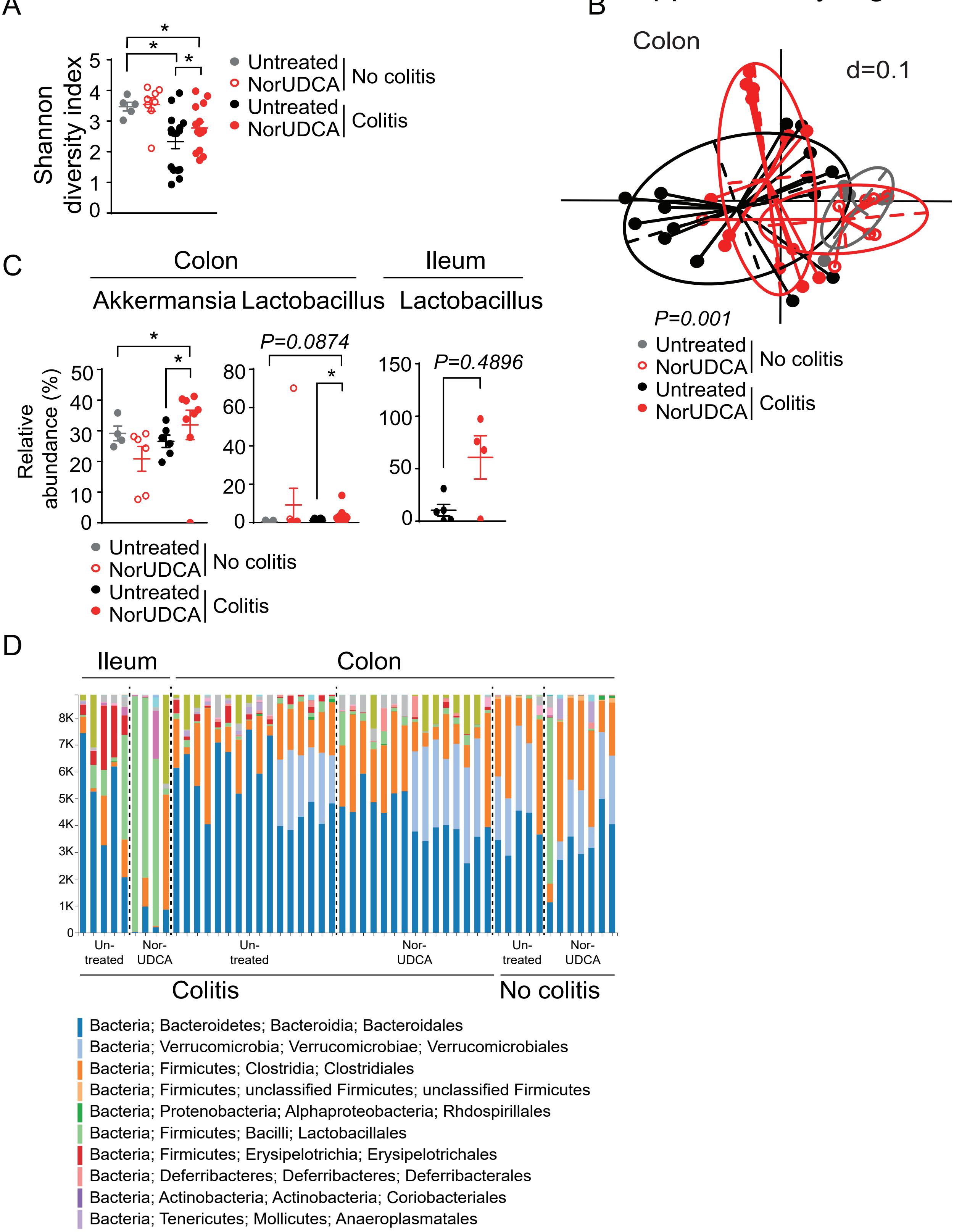

Supplementary Fig. 2

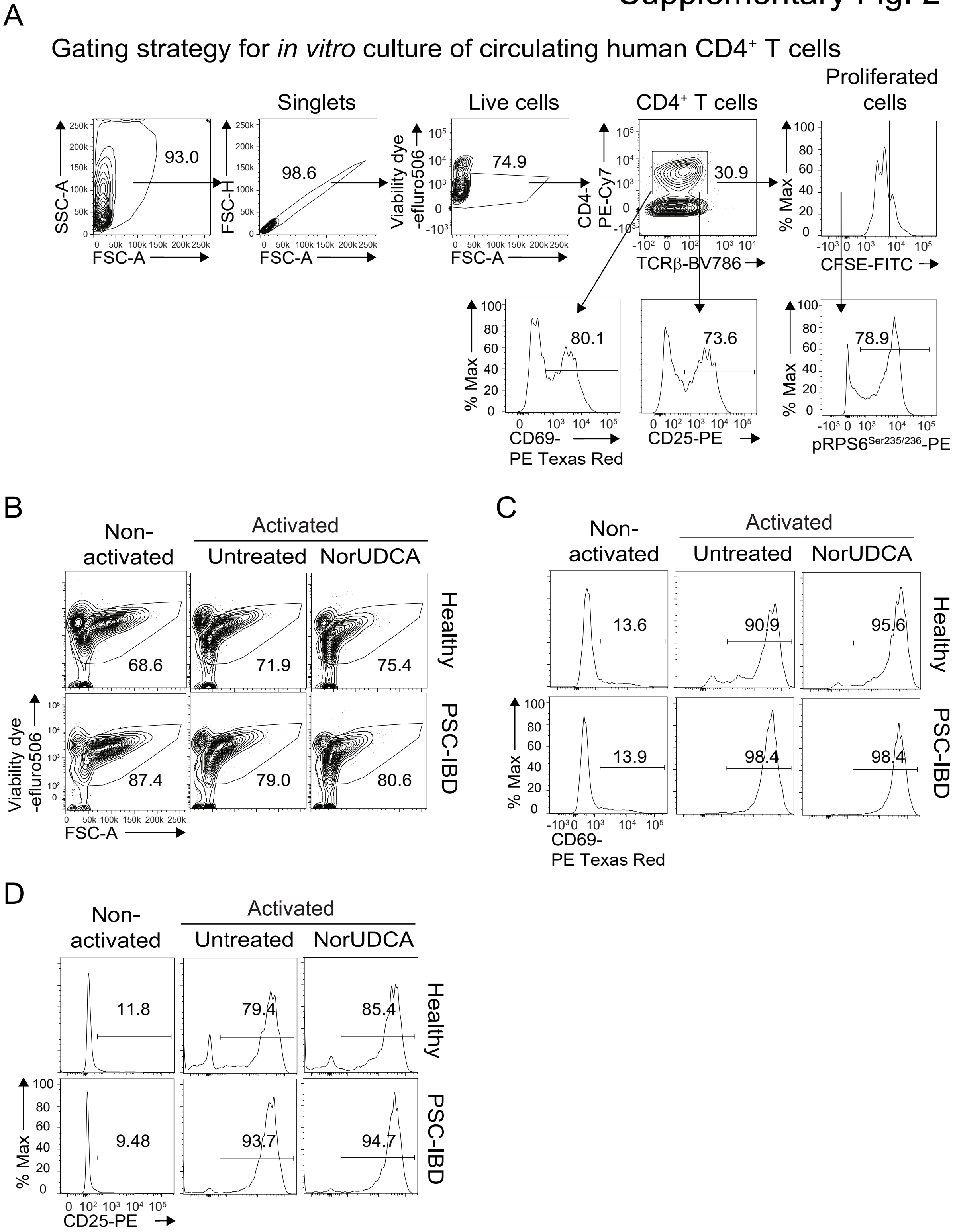
