## Supplementary material and methods for "24-Nor-ursodeoxycholic acid counteracts T_H_17/Treg imbalance and ameliorates intestinal inflammation by restricting glutaminolysis in differentiating T_H_17 cells"

Or

Wilfried Ellmeier, Dr.rer.nat.
Institute of Immunology, Center for Pathophysiology, Infectiology and Immunology, Medical University of Vienna, Lazarettgasse 19, A-1090 Vienna, Austria.


### Supplementary Material and Methods

#### Adoptive $CD4^+$ T cell transfer colitis model

To induce colitis, $CD4^+$ $T_{Naïve}$ cells were sorted from C57BL/6N mice by FACS and 0.5x$10^6$ cells were adoptively transferred into each *Rag2*$^{-/-}$ recipient mouse which either receive standard chow or 0.5% (wt/wt) NorUDCA-supplemented diet upon transfer. At week6, mononuclear cells from tissues of T cell reconstituted *Rag2*$^{-/-}$ mice were isolated, single cell suspensions were prepared from spleen, or mesenteric lymph nodes by mechanical disruption and passing through 70-µM nylon filters. For intestinal tissues, small intestines and colons were removed, rinsed thoroughly to remove fecal contents and opened longitudinally. Small intestine tissues were incubated with 5mM EDTA at 37°C for 30min to remove epithelial cells and dissociated in digestion buffer (RPMI1640, 1mg/mL collagenase type II, 5%FBS) with continuously stirring at 37°C for 1h. Mononuclear cells were collected at the interface of 40%-80% Percoll gradient. Cells were then analyzed for intracellular expression of transcription factors and/or cytokines after boosting with phorbol 12-myrisate 13-acetate (PMA; 5nM) and ionomycin (1µM) for 3-4h in the presence of Golgistop (10µg/mL). Colon tissue were made into swiss rolls for histology. Experiments were approved by the Austrian Federal Ministry for Science and Research at Medical University of Vienna (BMBWF-66.009/0039-V/3b/2019) and (BMBWF-2020.0.193.049).

#### *In vitro* $T_H17$ cell differentiation assay

Primary mononuclear cells were isolated from peripheral lymph nodes and spleens of C57BL/6N male mice (bred in the mouse facility of MUV) and magnetically enriched by for $CD4^+T_{Naïve}$ cells using Miltenyi Naïve $CD4^+$ T cell isolation kit. The purity of the $CD4^+T_{Naïve}$ cells was assessed by flow cytometry and was determined to be >98%. $CD4^+T_{Naïve}$ cells were stimulated *in vitro* with plate-bound anti-CD3 (1µg/ml), anti-CD28 (3µg/ml) Ab, polarized with recombinant mouse (rm)IL6 (20ng/mL), rhTGFb1 (1ng/mL), anti-mouse IFNγ (10µg/mL) and expanded in RPMI 1640 medium containing 10% fetal calf serum, GlutaMAX (2mM, ThermoFisher), β-mercaptoethanol (50µM, ThermoFisher) and Penicillin-Streptomycin (ThermoFisher). To test the impact of NorUDCA and/or $\alpha$-ketoglutarate ($\alpha$-KG) on $T_H17$ cell differentiation, culture medium±Glutamine was supplemented with NorUDCA and/or $\alpha$-KG as indicated in the figure. Viability of the cells was monitored by a fixable viability dye eFluor506 (ThermoFisher). Cells were analyzed 3 days later by flow cytometry (LSRII Fortessa, BD Biosciences).

### Supplementary Figure legends

**Supplementary Fig. 1. NorUDCA impacts on gut microbiome composition.** (A) Shannon index of microbiome diversity. (B) NMDS plot illustrating the gut microbiome diversity. (C) Relative abundance of selected taxa. (D) Relative abundance of gut microbiome. 4 biologically independent animals were used per group during experiments. Quantitative data are presented as mean±SE. *P* values were calculated by one-way ANOVA corrected for multiple comparisons with Dunnett test. *=$P$<0.05.

**Supplementary Fig. 2. NorUDCA does not affect human CD4+ T cell viability and activation.** (A) Gating strategy for flow cytometry analysis of *in vitro* culture of circulating human CD4+ T cells. (B) Viability profile of circulating CD4+ T cells from peripheral blood of healthy volunteer and PSC-IBD patients treated as indicated for 3 days. (C, D) Expression of CD25 and CD69 on live singlet CD4+ T cells of healthy volunteer and PSC-IBD patient are shown. Data in (B, C, D) are representative of 3 independent experiments. *P* values were calculated by one-way ANOVA corrected with Tukey post-hoc test. PSC, primary sclerosing cholangitis; Rapa, Rapamycin.

**Supplementary Table 1. Antibodies used in the study.**

| **Antigen** | **Clone** | **Company** | **Application** |
|---|---|---|---|
| CD4 | RM4-5 | Biolegend | F.C. |
| CD8a | 53-6.7 | Biolegend | F.C. |
| CD45R | B220; RA3-6B2 | Biolegend | F.C. |
| Gr1 | RB6-8C5 | Biolegend | F.C. |
| Ter-119 | TER-119 | Biolegend | F.C. |
| NK1.1 | PK136 | Biolegend | F.C. |
| CD25 | PC61 | Biolegend | F.C. |
| CD69 | H1.2F3 | Biolegend | F.C. |
| CD62L | MEL-14 | Biolegend | F.C. |
| CD8a | SK1 | Biolegend | F.C. |
| CD4 | SK3 | Biolegend | F.C. |
| CD69 | FN50 | Biolegend | F.C. |
| CD25 | BC96 | Biolegend | F.C. |
| TCRa/b | IP26 | Biolegend | F.C. |
| INFγ | XMG1.2 | BD Biosciences | F.C. |
| TNFα | MP6-XT22 | BD Biosciences | F.C. |
| CD3 | 145-2C11 | BD Biosciences | F.C. |
| CD28 | 37.51 | BD Biosciences | F.C. |
| CD3 | OKT3 | BD Biosciences | F.C. |
| CD28 | CD28.2 | BD Biosciences | F.C. |
| GLUT1 | EPR3915 | Abcam | F.C. |
| CD3 | SP7 | Novus | I.F. |
| Ecadherin | 4A2 | Abcam | I.F. |
| MPO | Polyclonal | Abcam | I.F. |
| CD44 | IM7 | Biolegend | F.C. |
| CD19 | 6D5 | Biolegend | F.C. |
| CD90.2 | 30-H12 | Biolegend | F.C. |
| FOXP3 | FJK-16s | Thermofisher | F.C. |
| pRPS6$^{S235/236}$ | D57.2.2E | Cell Signaling | F.C. |

| | | | |
|---|---|---|---|
| | | Technology | |
| pRPS6$^{S240/244}$ | D68F8 | Cell Signaling Technology | F.C. |

Table showing the antibodies used in this study. F.C., Flow Cytometry; I.F., Immunofluorescence.
